# Targeting anterior cingulate cortex with tDCS: an overview and analysis of electric field magnitude

**DOI:** 10.1101/2025.02.25.640124

**Authors:** U. Konstantinović, M. Stanković

## Abstract

The anterior cingulate cortex (ACC) is a brain region with a key role in various cognitive, emotional, and sensory processes. Given its extensive functional repertoire, modulating ACC activity to enhance cognitive functioning and alleviate symptoms of certain clinical conditions holds great potential. Over the past decades, transcranial direct current stimulation (tDCS) has gained popularity due to its ability to non-invasively affect the cortical excitability. Nevertheless, using tDCS to target regions located beneath superficial cortical areas, such as the ACC, could pose a challenge due to the unpredictable distribution of the electric field (E-field). To systematize the current state of evidence regarding the use of tDCS to modulate ACC activity, we conducted a systematic review focusing on analyzing the E-field distribution across the brain and its magnitude within the ACC as the region of interest. Our goal was to review the stimulation parameters used thus far and examine whether the E-field characteristics are linked to the observed effects. After the literature search and study selection, 14 studies were included in the review. Most of the studies were sham-controlled single-session experiments aiming to modulate specific cognitive processes in healthy adults. The most prominent process of interest was cognitive control, and most of the included studies observed behavioural effects. Our results show that cathodal stimulation led to significant results more frequently, regardless of E-field magnitude in ACC or specificity of E-field distribution throughout the brain. When it comes to anodal tDCS, more focal stimulation of the ACC was associated with a higher frequency of significant effects. We discuss these findings considering diverse methodological designs, the (un)specificity of ACC targeting, and other factors that could contribute to the variability of tDCS effects across studies.

## Introduction

Anterior cingulate cortex (ACC) is a brain region that is crucial for various aspects of human functioning. Generally, ACC is involved in cognitive and emotional functions, with particularly significant role in conflict monitoring and adaptation (Botvinick et al., 2001; Botvinick et al., 2004; Carter & Van Veen, 2007), pain processing (Fuchs et al., 2014; Xiao et al., 2021), decision making (Woo et al., 2022), error detection (Carter et al., 1998), response selection (Braver, 2001; Turken & Swick, 1999), addictive behavior (Zhao et al., 2021), risk-taking (Brown & Braver, 2008; Pei et al., 2020), emotion regulation (Giuliani et al., 2011), etc.

During the last two decades, techniques of non-invasive brain stimulation (NIBS) have emerged as promising tools for neuromodulation of many aspects of human behavior. Transcranial direct current stimulation (tDCS) is a NIBS technique that may affect cortical excitability of the stimulated cortical area. It is applied by setting up positive and negative electrode(s) on the participant”s scalp to form a constant, weak-current electric field throughout their head (Nitsche et al., 2008; Nitsche & Paulus, 2000). Even though tDCS has been successful in modulating diverse cognitive processes, targeting specific brain regions focally remains a challenge.

With its plethoric functional repertoire (Rolls, 2019; Stevens et al., 2011) ACC can be considered as a promising stimulation site for targeting a wide range of cognitive functions and various psychological states. On the other hand, anatomically, ACC is a part of cingulum cortex, which is wrapped around corpus callosum and lays in the longitudinal fissure, beyond the prefrontal cortex. Thus, due to its anatomical position, precise targeting of the ACC via tDCS is a challenge in the current state of art in the field. Using E-field distribution modeling is a fruitful approach to create an optimal stimulation protocol and make correct methodological decisions regarding specific research hypothesis.

To create an optimal stimulation protocol to reach the certain region of interest (ROI), stimulation parameters like current intensity, position, shape and polarity of the electrodes may be varied. This, in turn ultimately affects the distribution of the electric field. Thus, the electric field modeling approach creates an opportunity to simulate the electric field distributions induced by different electrode montages and tDCS protocols and test the electric field magnitude in the predefined ROI. The electric field modeling simulations may be conducted via modeling softwares: e.g. SIMNIBS (Simulation of Non-invasive Brain Stimulation (Saturnino et al., 2019, 2021; Thielscher et al., 2015) and ROAST (Realistic Volumetric-Approach-Based Simulator for Transcranial Electric Stimulation)(Huang et al., 2019). To fully explore the potential of targeting the ACC with tDCS, this systematic review aims to summarize findings from all available tDCS studies using the ACC as a stimulation target. Our review will focus on the effects of ACC-targeted tDCS on cognitive and psychological processes. We intend to summarize the existing montages and analyze the effects in terms of E-field distribution across the brain and E-field magnitude within the ACC as the region of interest (ROI). Since ACC is structurally divided into five sub-regions (pregenual, subgenual, dorsal, caudal, rostral), we focused on dorsal ACC (dACC), as a ROI since it is most frequently associated with cognitive and emotional processing (Bush et al., 2000).

## Methods

The review was conducted following the guidelines from 2020 Preferred Reporting Items for Systematic Review (PRISMA) (Page et al., 2021).

### Selection criteria

The primary inclusion criterion for study selection was the use of tDCS to modulate the activity of the ACC. We included studies on either healthy participants or participants presenting with clinical symptoms that do not affect neuroanatomy. Thus, we excluded studies on participants with diagnoses such as traumatic brain injury or stroke, since this could represent a confounding factor for accurate modeling of the E-field. We excluded studies with other Transcranial Electrical Stimulation techniques (e.g., Transcranial Alternating Current Stimulation) since standard modelling approaches for those techniques could be unreliable due to increased complexity in accurately estimating E-field distributions. Additionally, we included studies of any methodological design, both sham-controlled and non-controlled studies. Finally, we included studies published in English language in peer-reviewed journals.

### Search strategy

We performed a search in PubMed using the following search string: (“transcranial direct current stimulation”[Title/Abstract] OR tDCS[Title/Abstract]) AND (“anterior cingulate cortex”[Title/Abstract] OR ACC[Title/Abstract]). To identify potentially missed publications, the authors manually searched Google Scholar and references of the previously identified publications.

We used the Rayyan automation tool for abstracts and full text screening. U.K and M.S independently screened the abstracts in blind mode. Conflicts were resolved with the help of a third researcher (J.B.).

### Data extraction and E-field strength calculations

The search was conducted on January 13, 2025. We retrieved the following data for each study: sample size, participant population (i.e., healthy or clinical), study design, study protocol, stimulation duration, stimulation intensity, electrode placement, electrode size and shape, cognitive function/symptoms assessed, outcome measures, and main results.

Based on the parameters for electrode placement, electrode shape and size, and simulation intensity, we calculated E-field for each utilized montage. We used the SIMNIBS software (Saturnino et al., 2019, 2021; Thielscher et al., 2015) to calculate the E-field intensities. The simulations were conducted on the example dataset m2m_MNI152 in the SIMNIBS package. The region of interest (ROI) was defined using MNI coordinates for the dorsal ACC (−5; 14; 42 for left hemisphere and 5; 14; 42 for right hemisphere) (Zhou et al., 2016), with a sphere radius of 10 mm. To evaluate the mean electric field magnitude in a grey matter ROI, we first utilized the *mni2subject_coords* function in MATLAB environment for transforming the MNI coordinates to the average head model (m2m_MNI152) space after loading the simulation output. Conductivity values for various tissue types, as determined in prior studies, were utilized: σskin at 0.465 S/m, σbone at 0.01 S/m, σcerebrospinal fluid at 1.654 S/m, σgray matter at 0.275 S/m, and σwhite matter at 0.126 S/m (Windhoff et al., 2013). The average field strength in this region was then calculated, with higher values indicating stronger electric fields.

## Results

### Study selection

The literature search yielded 93 studies for screening. Figure 1 illustrates the study selection process. Ultimately, 14 studies met the inclusion criteria.

**Figure 1.**
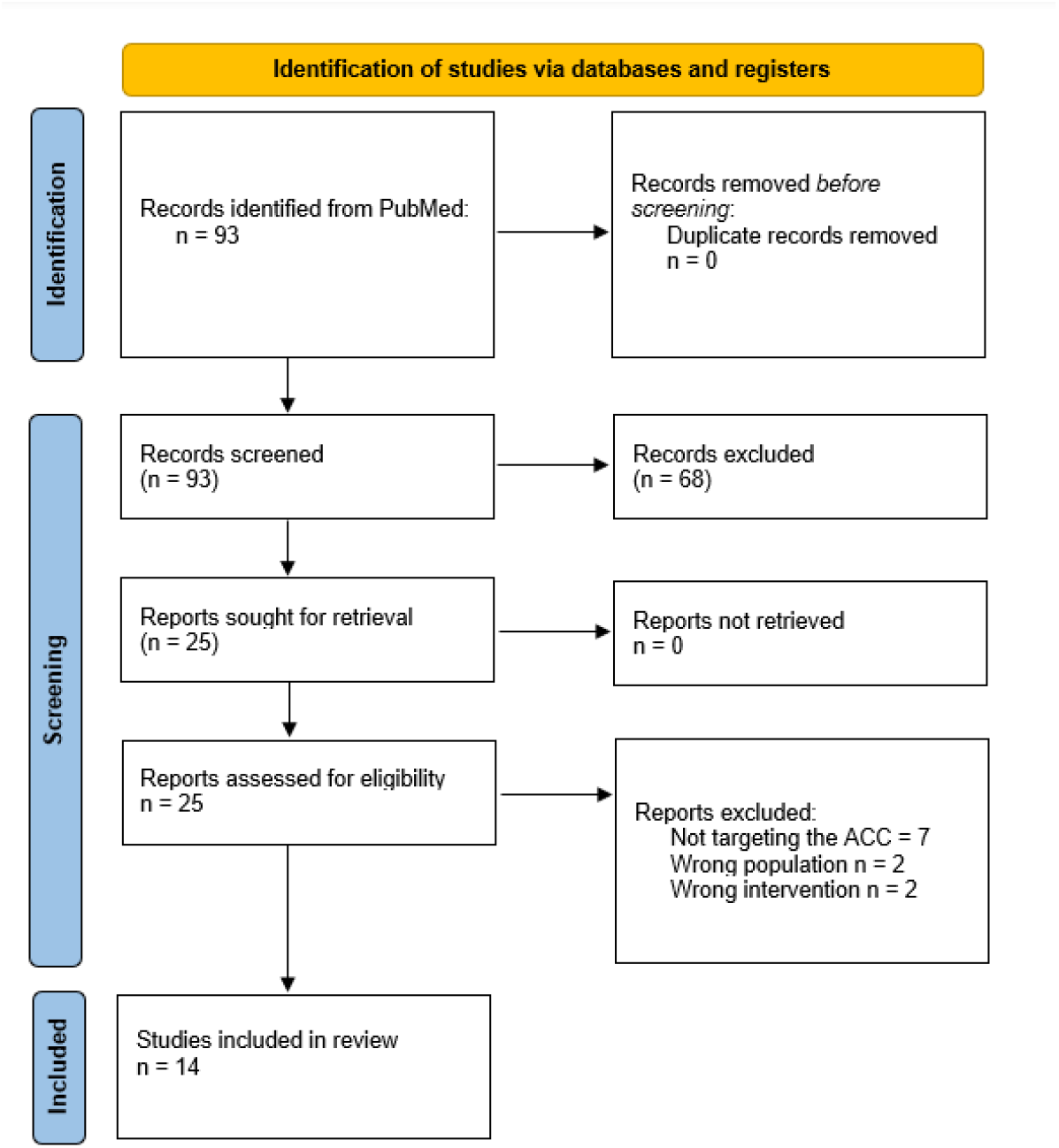
A PRISMA chart showing the study selection process

### The study characteristics

Table 1 synthesizes all the features extracted from the studies in this review. Regarding study design, both between-subject and within-subject design were equally represented (43 %) in the observed studies while mixed design was employed in 14% of the studies. Study population of interest consisted mostly of heathy adults (71%), with four studies having clinical population as a sample. The clinical conditions they focused on included headache, low back pain, HIV, and comorbid depression and anxiety.

**Table 1:**
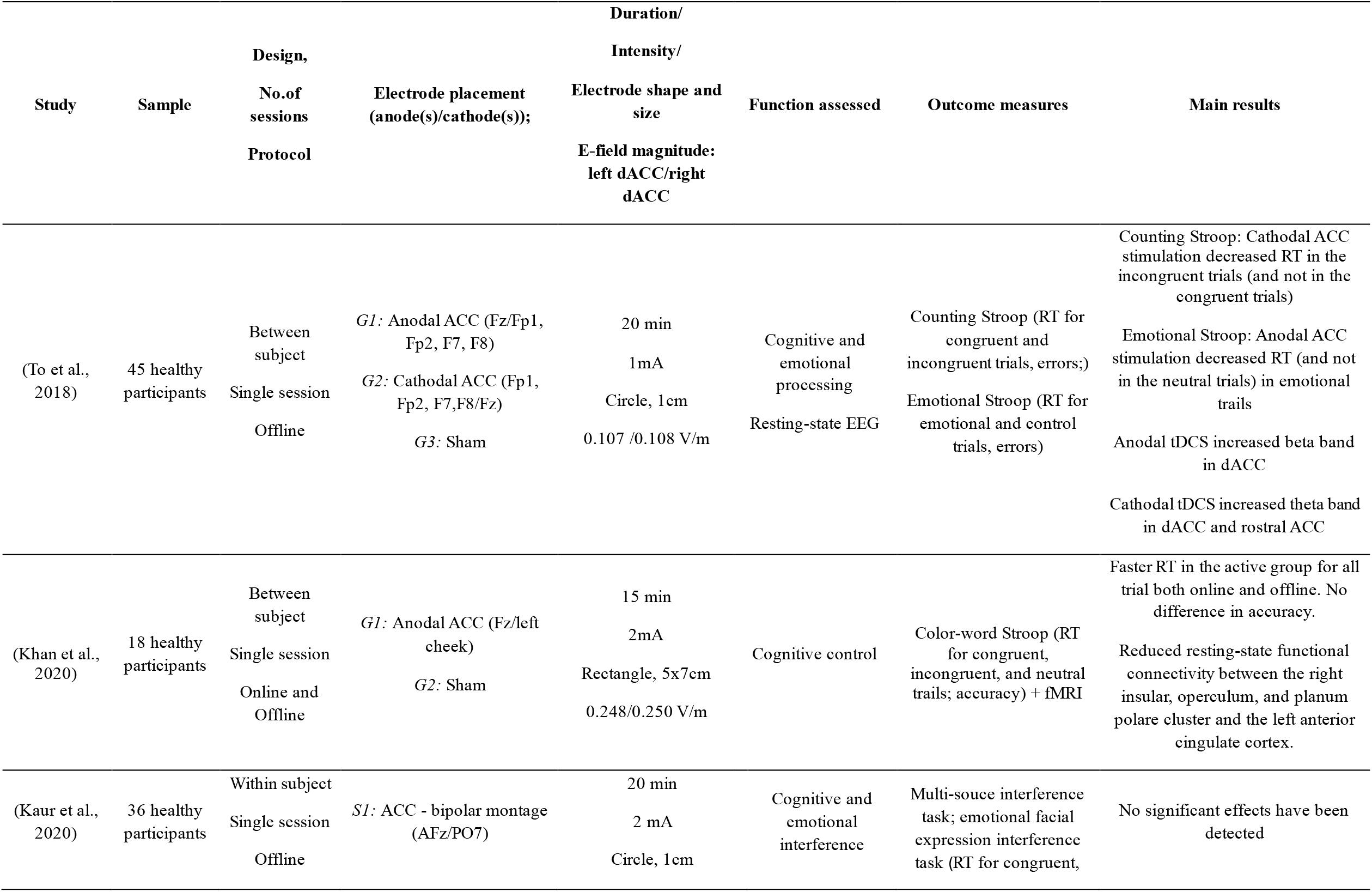
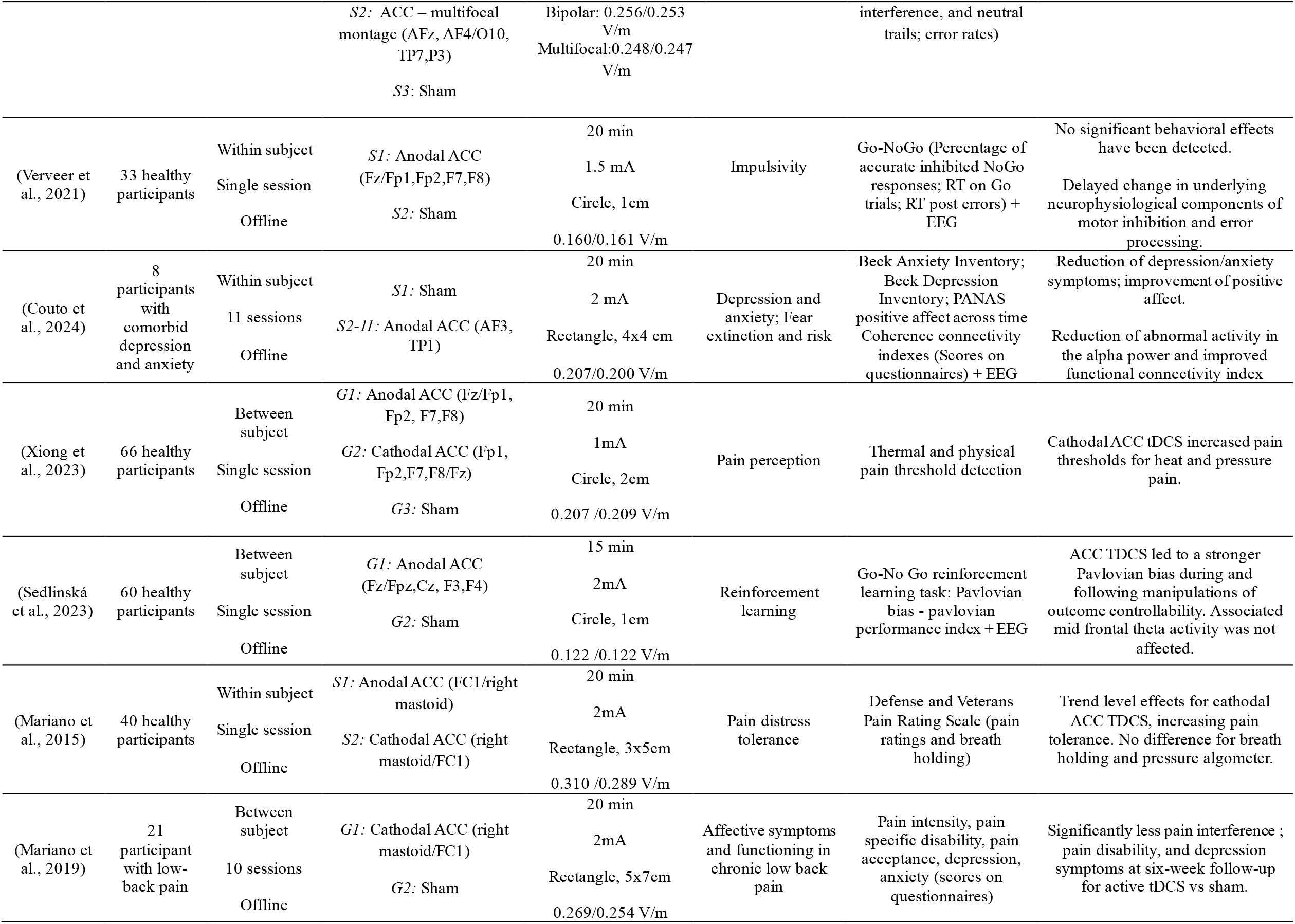
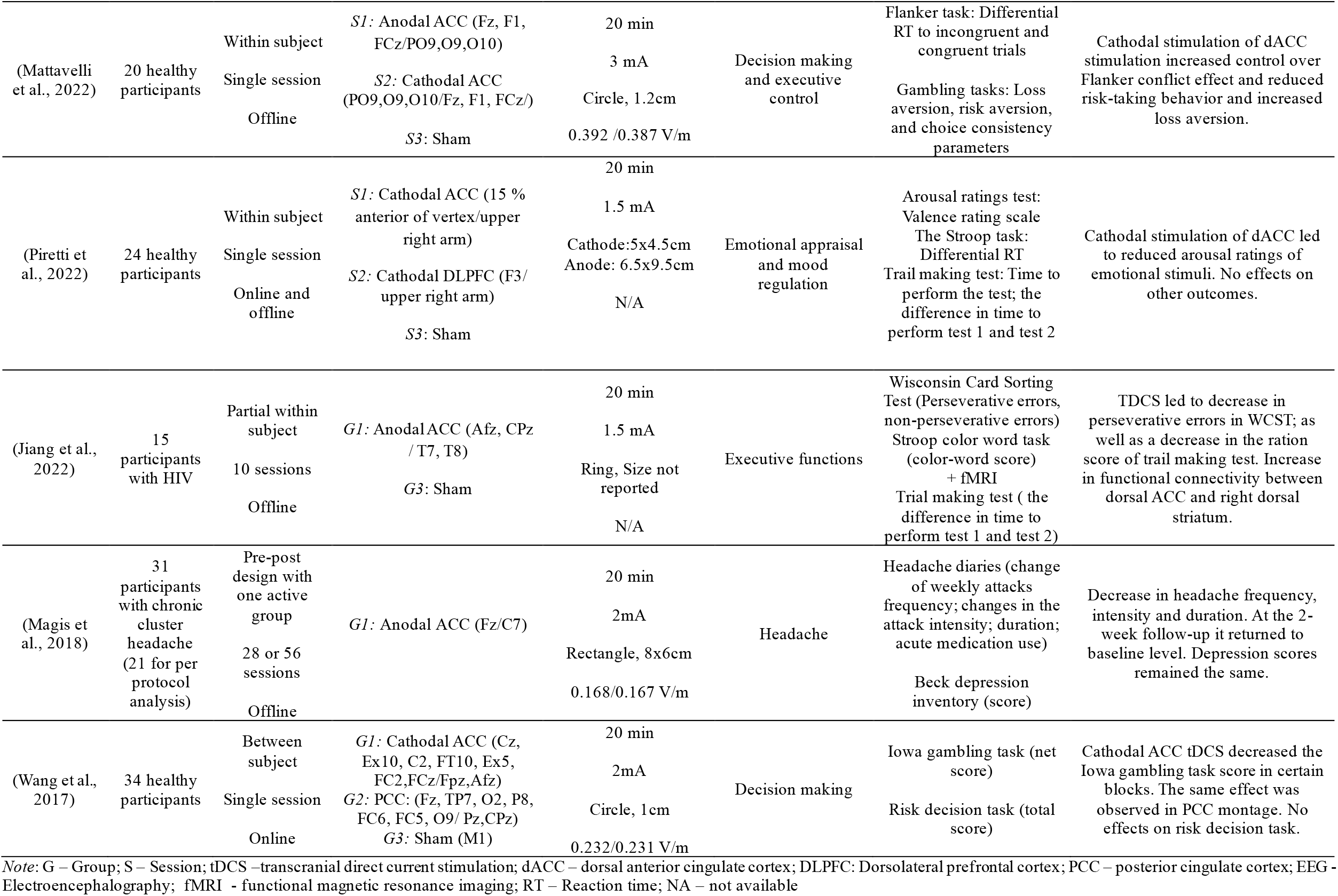
Characteristics and main results of studies that used tDCS techniques to modulate ACC activity

Majority of studies (71%) were designed as single-session studies as opposed to multi-session design. The bipolar and multifocal electrode montages to target ACC are almost equally represented in all studies, with multifocal montages being slightly more frequent (57%). Not all the studies were sham-controlled, since 2 of 14 didn”t have sham condition. The most common position for the stimulation electrode was Fz (43%). Specifically, five studies used Fz as a stimulation site in a bipolar montage, while one study included an Fz electrode in a multifocal setup. Another frequent stimulation electrode position, used in three studes was AFz. The most frequent electrode placement used in multiple (n = 3) studies was: Fz / Fp1, Fp2, F7, F8; (To et al., 2018; Verveer et al., 2021; Xiong et al., 2023). Seven studies aimed to affect the excitability of the ACC with anodal stimulation; conversely, three studies attempted the opposite, i.e., to modulate the ACC with cathodal stimulation. The remaining four studies utilized both anodal and cathodal stimulation of the ACC. Stimulation intensity range varied throughout the studies from 1 to 3 mA with the majority that used 2 mA (64%). The duration of tDCS treatment was 20 min in 12 of the 14 studies, while in the remaining two, tDCS was applied for 15 min. The electrode size and shape were mostly a 1 cm diameter circle in studies using a multifocal montage, while the studies with bipolar montages mostly used rectangular electrodes with dimensions ranging from 4 to 7 cm. According to ACC involvement in multiple cognitive and emotional functions, observed studies were focused on effects of tDCS on various behavioral functions. The substantial number of studies are in the cognitive domain, and they were most frequently focused on executive functioning, decision making, cognitive control and interference processing. There was a certain number of studies focusing on the effects of tDCS on pain perception and processing, while a few studies focused on emotional processing. Regarding the outcome measure, different modifications of Stroop task were most commonly used (in four studies) with mean reaction time (RT) for every category of trials as a most frequently used score. Other cognitive tasks and assessment scales were each utilized in only a single study, with the exception of the Go/No-Go task.

### The E-field magnitude in dACC and effects of tDCS

Table 1 illustrates that the results of almost all studies that involved a cathodal tDCS condition in their design showed significant effects on at least one outcome measure. Out of the seven studies where cathodal stimulation was applied, the results of five showed boosting of the critical function studied. One studyobtained function reduction, and one study showed a positive significant effect on a trend level. In total, of 17 outcome measures, cathodal stimulation had significant effect on in 10 cases. On the other hand, the results of studies with anodal stimulation are more conflicting. In three of the nine studies, researchers did not report any significant effect of anodal tDCS of ACC. Nevertheless, in five studies, significant effects were observed, with one that impaired and four enhanced the function of interest. In summary, the anodal tDCS over ACC had significant effects in 9 out of 24 cases.

The calculated E-field magnitude in dACC ranged between 0.107 V/m to 0.392 V/m, with the average value of 0.223 V/m across all studies. For two studies E-field magnitude could not be calculated due to lack of detailed method section reporting. For cathodal stimulation, studies with electrode montages ranging from highest (0.39 V/m), to the lowest (0.11 V/m) dACC E-field magnitude showed significant behavioral effects. Notably, the only study applying cathodal stimulation with zero effects for all outcome measures was using the tDCS protocol that induced the second highest E-field magnitude in dACC. Additionally, the obtained results of two studies, from which we could not calculate the E-field magnitude due to a lack of reported information, shows that only one-third of the effects are significant. For anodal stimulation, results are showing zero behavioral effects of studies that applied montages with highest E-field magnitude in dACC. Specifically, significant effects are obtained only in studies with electrode montages that induced medium or lower magnitude of E-field in dACC – around 0.20 m/V or bellow.

### (Un)specificity of tDCS effects on ACC

Figure 2 presents the E-field distributions as results of SIMNIBS simulations outputs for each study observed in this review. In all studies E-field was predominantly focused on medial lateral prefrontal cortex. However, it can be observed that E-field was distributed differently throughout the brain in different studies, leading the stimulation more or less specific for the ROI defined.

**Figure 2.**
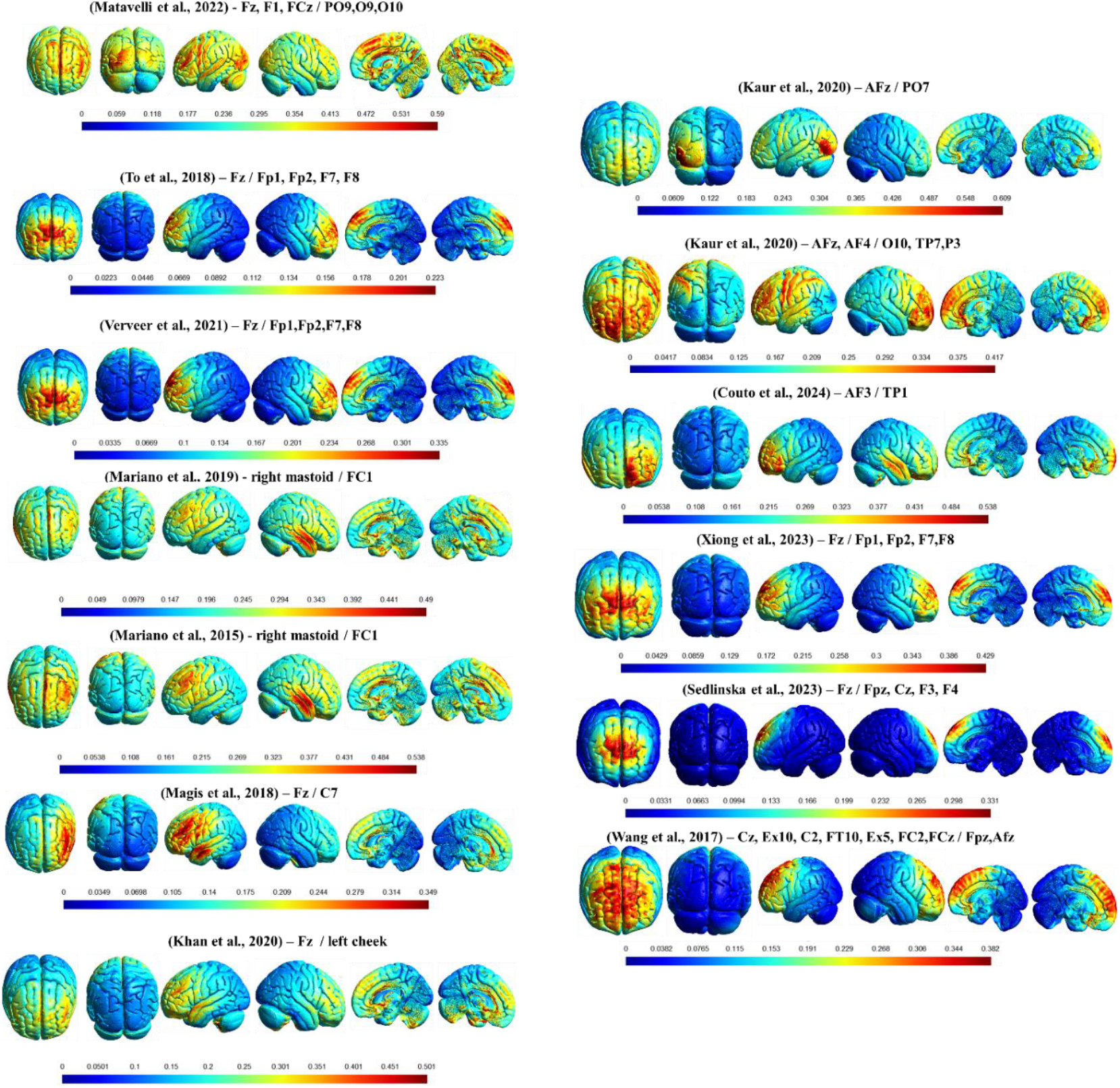
The output of SIMNIBS simulations for every ACC tDCS montage in all observed studies in the systematic review. Electric field distribution is presented in six different angles and planes (frontal, occipital, left, right, sagittal left, sagittal right) for every observed tDCS montage. The color map scale for every montage is in V/m.

Inspection of Figure 2 indicates that the behavioral effects of cathodal ACC stimulation are not dependent on the specificity of E-field distribution throughout the brain. Some of this studies (Mariano et al., 2019; Mattavelli et al., 2022) used electrode montages that induced broad E-field and yielded significant enhancing effects on multiple behavioral outcomes. Another study with a broad E-field montage produced a significant effect as attenuating decision making, but had zero effect on risk control (Wang et al., 2017). Still, studies with more focal montages (To et al., 2018; Xiong et al., 2023) resulted in positive significant behavioral effects. On the other hand, the only study with non-significant effects (Mariano et al., 2015) had the same montage as the aforementioned study of the same authors (Mariano et al., 2019), but with smaller electrodes resulting in a less distributed E-field.

Conversely, studies with anodal tDCS on ACC are more frequently yielding significant behavioral effects with electrode montages that induce more focal E-field distribution. Results of four studies (Couto et al., 2024; Khan et al., 2020; Magis et al., 2018; To et al., 2018) showed positive significant behavioral effects. Additionally, another study (Sedlinská et al., 2023) with montage producing focal E-field distribution resulted in a significant effect but in the opposite direction of the hypotheses. However, one study (Verveer et al., 2021) with electrode positioning that induced focal E-field distribution didn”t show any significant effect. On the other hand, studies that used electrode montage with broader E-field distribution (Kaur et al., 2020; Mariano et al., 2015; Mattavelli et al., 2022; Xiong et al., 2023) didn”t observe any effects of anodal stimulation, regardless of the calculated E-field magnitude in dACC.

## Discussion

### Heterogenous methodology

Although there is a proliferation of studies with tDCS in the field (Sun et al., 2022), number of studies with tDCS over ACC is still limited to a dozen papers. It is observed that the majority of studies in which ACC was stimulated significantly differed from one another in terms of methodological aspects. The electrode size, shape, and positioning varied from study to study, leading to different E-field distribution in almost every protocol, which ultimately affected the comparability of the results obtained. However, the montage with electrodes positioned at Fz / Fp1, Fp2, F7, F8 used in multiple studies (To et al., 2018; Verveer et al., 2021; Xiong et al., 2023) with different current intensity, yielded inconsistent results. An additional source of uncontrolled variability which may hinder comprehensive evaluation of effects ACC-targeted tDCS is the fact that almost every study focused on different function, assessed by various of tasks and measurement scales. Thus, distinguishing the specific tDCS effects over ACC is a complex challenge since it seems to be inevitable that same stimulation dose leads to the significant effect on one outcome measure but not on the other. Consequently, there is a need for new studies with more uniform stimulation parameters and outcome measures to obtain more reliable evaluation of the effects. Since there is an ongoing replication crisis in neuroscience (Button et al., 2013), the more homogenous methodology would enable potential replication experiments which would strengthen the empirical evidence of ACC tDCS effects.

### E-field magnitude – the more the merrier?

By comparing the behavioral effects of studies with tDCS over ACC and results of dACC E-field magnitude calculations we may raise a question of potential impact of fine-tuning differences in E-field magnitude on ultimate behavioral effects. Does a higher E-field magnitude per se yields stronger effects? Results of our review are not completely conclusive for adequate response to this question. Studies that showed significant effects of cathodal stimulation are not directly dependent on E-field magnitude in dACC – protocols with the wide range (0.11-0.39 V/m) of E-field magnitude leaded to significant effects. On the other hand, for anodal tDCS protocols, studies with the highest E-field magnitude leaded to non-significant effects, while studies with lower and moderate values of E-field magnitude in dACC yielded significant effects. In line with this information, we can draw a conclusion that, at least for anodal tDCS, higher E-field magnitude doesn”t independently assume larger and significant behavioral effects of stimulation.

### Specificity of the effects of ACC stimulation

Impact of the E-field magnitude in dACC on behavioral effets of ACC tDCS could be better understood in relation to the E-field distribution. While applying tDCS to predetermined ROI, it should be considered that the stimulation is introducing a certain level of both signal and noise to the system (Miniussi et al., 2013). Signal represents specific effects of stimulation on ROI, while noise can be defined as its unspecific effects including stimulation of other brain areas, perception of cognitive task that has been used, participant”s current state, etc. The ultimate effects of stimulation depend on signal/noise balance. Effectively targeting the ACC with tDCS presents methodological limitations, as researchers must decide which additional, more surface-level brain regions will also be stimulated due to the ACC”s deeper location in the brain. Results in our review indicate that cathodal stimulation was effective regardless of broadness/focality of E-field distribution. On the other hand, anodal stimulation yielded significant effect only in studies with electrode montages that produced more focal E-field mainly within the medial prefrontal cortex (MPFC). This implies that anodal stimulation may produce effects when the E-field is specifically focused in MPFC, with minimization of stimulation of other regions.

Additionally, a limitation of tDCS studies targeting the ACC is the propagation of unspecific effects on the activity of other brain areas functionally related to the outcome measures. For example, results of the studies that assessed effects of ACC tDCS on cognitive control / inhibiton outcome measures, like Stroop task (Kaur et al., 2020; Khan et al., 2020; To et al., 2018) may be affected by unspecific stimulation of presupplementary motor area (pre-SMA) or lateral prefrontal cortex areas which are crucial nodes for performance in cognitive inhibition tasks (Hsu et al., 2011; Jacobson et al., 2012). Thus, it may be difficult to disentangle if the effect is induced by specific (ACC) or unspecific (pre-SMA, lateral prefrontal cortex) stimulation, if a study relies only on evaluation of behavioral effects of ACC tDCS on the mentioned measures. This limitation can be overcome by using additional physiological readout measures like source localization EEG (To et al., 2018) or resting state fMRI before and after the stimulation (Khan et al., 2020), to more accurately evaluate specific ACC stimulation effects.

## Conclusion

To conclude, in this review, we synthesized the results of tDCS studies targeting ACC with a particular focus on relation of E-field magnitude and distribution and behavioral effects obtained in the literature. Cathodal stimulation was more frequently inducing significant effects, regardless of E-field magnitude in dACC or specificity of E-field distribution throughout the brain. On the other hand, anodal tDCS appears to induce significant effects in studies that used more focal electrode positioning, with moderate E-field magnitude in dACC.

